# Associating with kin selects against disease tolerance

**DOI:** 10.1101/2023.09.04.555776

**Authors:** Jason Cosens Walsman, Madalyn Lambe, Jessica F Stephenson

**Affiliations:** University of Pittsburgh

## Abstract

Behavioural and physiological immunity are key to slowing epidemic spread. Here, we explore the evolutionary and epidemic consequences of their different costs for the evolution of tolerance vs. resistance: behavioural resistance affects social cohesion, with associated group-level costs, while physiological resistance costs should accrue to the individual. Further, the transmission-reduction benefits of resistance accrue differently to susceptible hosts and those already infected; infected hosts only benefit indirectly, by reducing transmission to kin. We therefore model the coevolution of transmission-reducing defences expressed in susceptible hosts with those expressed in infected hosts, as a function of kin association, and analyse the effect on population-level outcomes. Using parameter values for guppies, *Poecilia reticulata*, and their gyrodactylid parasites, we find that: 1) either susceptible or infected hosts should invest heavily in preventing infection, but not both; 2) kin association drives investment in physiological resistance more strongly than in behavioural resistance; 3) even weak levels of kin association can favour altruistic infected hosts that invest heavily in resistance (vs. selfish tolerance), eliminating the disease. Overall, our finding that weak kin association affects the coevolution of infected and susceptible investment in both behavioural and physiological immunity suggests that kin selection may affect disease dynamics across systems.

## Introduction

Hosts employ multiple defences to prevent disease transmission [1-3] and these defences differ in their costs and benefits. For example, transmission-reducing, asocial behaviour (an example of behavioural immunity) is likely to have costs in terms of reduced social benefits [4, 5], experienced by the focal, asocial individual and possibly also by non-focal individuals missing out on the benefits of social contact [e.g., smaller groups hurt fitness of various social mammals: 6]. Physiological immune investment (physiological immunity at times referred to as “tissue-specific” [7]), on the other hand, acts within the body of an individual host to prevent transmission and often carries internal, physiological costs borne only by the focal individual expressing the immunity [8-11]. Costs of behavioural or physiological immunity can be borne by susceptible hosts, or infected hosts, e.g., costly self-isolation (behavioural immunity) or costly reduction of parasite shedding (physiological immunity). However, the transmission-reduction fitness benefits of immunity accrue differently to susceptible hosts and those already infected: while susceptible hosts directly benefit from transmission-blocking immunity, infected hosts only benefit indirectly, by reducing transmission to kin.

This topic touches on critical, ongoing questions of host evolution of resistance vs. tolerance. While much previous work has focused on physiological resistance or tolerance, a growing body of work emphasizes the importance of behavioural resistance or behavioural tolerance [7, 12]. While “resistance” and “tolerance” vary significantly in definition, we use a broad definition in which a more resistant host genotype directly lowers parasite fitness while a more tolerant host genotype does not lower parasite fitness but has higher fitness given the same infection [following the definition of 13]. So susceptible hosts that prevent infection are resistant, behaviourally or physiologically. Infected hosts that prevent onward transmission are also resistant. Infected hosts that do not make the costly investment to prevent onward transmission are tolerant, i.e., they have higher fitness given the same infection. Thus, we consider infected hosts that face a tolerance-resistance trade-off (empirically demonstrated for physiological immunity [14-17] and with some support in behavioural immunity [12, 18, 19]); specifically, we consider a tolerance-resistance trade-off in which infected resistance does not improve health as much as tolerance does [empirically supported in some systems; 15, 18, 19] so that infected resistance only makes sense by benefiting kin.

In this context, infected hosts should only invest in costly resistance if kin selection is very important relative to selection on direct fitness [20, 21]. This has been empirically observed as self-isolation (behavioural resistance) in eusocial insects, in which kin association is very frequent and thus kin selection should be extremely important [18, 19]. However, investigation of active self-isolation is lacking in other systems [21], potentially because it can be difficult to disentangle when self-isolation is beneficial for the infected host’s direct fitness [22]. For physiological resistance, some previous modelling has shown that kin association can favour transmission-blocking by infected hosts [23; their model has individual costs]. An extreme form of such physiological resistance, suicide by infected hosts, evolved in clonal bacterial populations with spatially-driven kin association [24]. But many studies, theoretical and empirical, have assumed kin association is usually too weak to matter for evolution of altruistic resistance by infected hosts [we use “altruistic” to refer to a trait that reduces direct fitness and increases the fitness of other individuals, following Hamilton: 25]. Instead, much previous work has either ignored evolution of infected resistance or considered only selection on direct fitness for infected hosts. In nature, however, kin association falls across a broad range, e.g., with estimates of within-group relatedness ranging from nearly 0 to 0.78 across vertebrate taxa [26-29]. So how strong does kin association need to be for altruistic infected resistance to matter?

This question of infected resistance cannot be adequately addressed without considering the feedbacks between the evolution of infected resistance and susceptible resistance. If infected hosts evolve more or less resistance (behavioural or physiological), then susceptible hosts may as well. Such differential investment in resistance has received particular empirical attention for behavioural immunity. Infected hosts have expressed lower gregariousness [higher behavioural resistance; 18, 19, 30] or higher gregariousness [31, 32] than susceptible hosts. Resistance in one class will feed back on selection on the other class’s trait, requiring a model to tease out the feedbacks. Little of the existing intuition and modelling have considered interactions between susceptible immune investment and infected immune investment, particularly in regards to kin selection [as noted by 23].

Lastly, how do these questions apply to the cases of behavioural and physiological resistance? Long-standing and sustained attention has been given to understanding the action and relative importance of behavioural and physiological immunity [1, 3, 33-37], particularly because they can govern transmission rate in similar ways. However, very little theoretical work has directly compared and contrasted the evolution of these defences [38]. We are unaware of any that consider kin selection, or the shared costs of behavioural resistance versus the individual costs of physiological resistance [33 uses shared costs of behaviour but without contrast to fixed]. Thus, kin selection may interact differently with the coevolution of resistance in the behavioural resistance case and the physiological resistance case.

We model coevolutionary outcomes for both cases across a range of kin association using parameter values relevant to a focal, empirical system. We use parameter values [39] from a guppy-worm system (*Poecilia reticulata*-*Gyrodactylus* spp.) with social transmission [40] and heritable variation in behavioural resistance governing contact [41, 42] and physiological resistance [43, 44]. There is some evidence for kin association in guppy populations. Juvenile guppies show some ability to detect kin in laboratory conditions [45] and wild populations vary in whether juveniles are disproportionately related to their shoalmates [27] but wild adults are not disproportionately related to their shoalmates [45, 46]. Further, uninfected and infected hosts can differ in their behavioural resistance [3] as infected guppies may seek contact [3, 47] while uninfected guppies may avoid contact [40, 48, 49], potentially reducing overall rates of social contact [49]. Thus, this focal system provides empirically-derived parameter values as well as empirical motivation for model ingredients like weak kin association and infection-class differences in behavioural resistance. To see a range of possible outcomes that may be broadly relevant to various systems, we consider a range of kin association from none to very strong. We compared and contrast results for behavioural and physiological resistance; for both models, selection for altruistic resistance in infected hosts can occur even when kin association is rare. This result has important implications for host populations as well as the simultaneous coevolution of resistance and tolerance.

## Methods

We model coevolution of resistance investment by susceptible and infected hosts with a differential equation model and evolutionary invasion analysis. For both behavioural or physiological resistance, evolution is represented in the trait *g*_*X, Y*_, which depends on genotype *Y* and infection status *X* (*S* for susceptible, *I* for infected, or *H*=*S*+*I*). Higher values of *g*_*X, Y*_ correspond to higher transmission rates given encounter between a susceptible and infected host, i.e., lower resistance. For the behavioural resistance model, *g*_*S, Y*_ represents the gregariousness of susceptible hosts of genotype *Y* and *g*_*I, Y*_ is the gregariousness infected hosts. Higher gregariousness generally leads to more social contacts for the focal individual (*C*_*XY*, Total_ is the total rate of social contacts for an individual of genotype *Y* and infection status *X* with the rest of the host population) as well as for non-focal individuals; more social contacts lead to lower death rate, e.g., living in groups helps defend against predation in many systems [50] including the focal guppy system [51] (predation death is *B*/*C*_*XY*, Total_ where *B* scales predation).

For the physiological resistance model, *g*_*S, Y*_ is the infectability of susceptible hosts of genotype *Y* and *g*_*I, Y*_ is the infectiousness of infected hosts; physiological resistance investment lowers *g*_*X, Y*_ and thus transmission but carries a mortality cost borne by the focal individual in terms of increased death rate, e.g., due to immunopathology [52] (immunopathology death is *B*/*g*_*X, Y*_). In both models, resistance reduces transmission rate (lower *g*_*X, Y*_) but carries a cost of increased mortality borne either by the focal individual (physiological resistance) or by the focal individual and non-focal individuals (behavioural resistance).

Resistance reduces the chance of transmission given encounter between an infected and uninfected individual. Each individual has a fixed *E* encounters per day which are divided into random encounters (*R*) with all genotypes at frequency-dependent rates and non-random, assortative encounters that are only with kin [1-R, following the classic model of 53]. Higher rates of random encounters correspond to lower values of kin association. For simplicity, we only consider kin as individuals of the same, clonal genotype. Thus, the function for encounters per time unit experienced by an individual of class *X*_Y_ with individuals of class *W*_Z_ is given by Eq. 1a. Note that the *W*_*Z*_ class may or may not be the same class as the *X*_*Y*_ class of the focal individual.

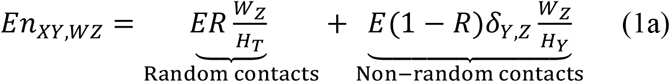

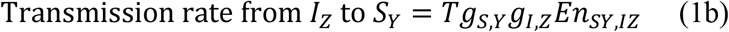

Here *δ*_Y, Z_ is the Kronecker delta (1 when *Y* = *Z* and 0 when *Y* ≠ *Z*); thus, individuals have random contacts with kin and non-kin but non-random contacts only occur between individuals of the same genotype (i.e., are kin). Transmission from infected hosts (e.g., of genotype *Z*) to susceptible hosts (e.g., of genotype *Y*) then occurs at a rate directly proportional to encounter rate (Eq. 1b). Transmission rate is also directly proportional to some transmissibility parameter (*T*) and the *g* parameters that depend on resistance of susceptible (*g*_*S,Y*_) and infected (*g*_*I,Z*_) hosts. In the behavioural resistance model, *g*_*X,Y*_ is gregariousness and can be thought of as the probability of accepting contact given encounter (ranging from 0 to 1), contributing to the joint probability of both individuals accepting contact gives the contact rate (*C*_*XY, WZ*_ = *g*_*X,Y*,_ *g*_*W,Z*_ *En*_*XY, WZ*_); thus, the gregariousness of another individual (*g*_*W,Z*_) influences the contact rate an individual experiences and thus the mortality cost of resistance investment is shared (e.g., higher *g*_*W,Z*_ can lead to higher *C*_*XY, WZ*_, thus higher *C*_*XY, Total*_, and thus lower death rate of *X, Y* individuals). In the physiological resistance model, these *g* parameters are simply susceptible infectability and infected infectiousness, respectively, and they simply scale transmission rate (also limited to 0-1 for comparability to behavioural resistance). Since each individual has a fixed *E* encounters per day, disease transmission is frequency-dependent for both models. Frequency-dependent transmission may be reasonable for our focal system [54] and is a necessary assumption or else different levels of kin association would artificially alter disease transmission, e.g., any rare genotype would suffer essentially zero infection in the case of all non-random, assortative encounters and density-dependent transmission. We embed this function for encounters and disease transmission within an SI model of host densities.

We model ecological dynamics of the susceptible host density, *S*_r_, and infected host density, *I*_r_, of a resident genotype, r. Hosts grow logistically with maximum fecundity *b* and sensitivity to crowding *q*; both births and crowding depend on the total density of the resident genotype: *H*_r_ = *S*_r_+*I*_r_ (Eq. 2a). Susceptible hosts die with some background mortality rate *d* and additional mortality reflecting the cost of resistance. Susceptible individuals are also lost to infection depending on transmissibility of infection, *T*, and resistance of susceptible (*g*_*S,r*_) and infected hosts (*g*_*I,r*_), and the rate of encounters with infected hosts (*En*_*S*r,*I*r_). Once infected, hosts suffer background mortality and additional mortality related to resistance investment; they also suffer additional mortality due to the virulence, *v*, of infection (Eq. 2b). These equations are very similar for the behavioural resistance model (Eq. 2a, b) and the physiological resistance model (Eq. 2c, d) except that the cost of resistance depends on resultant total contacts for behavioural resistance (written out for clarity in Eqs. 2a, b) but simply on resistance for physiological resistance.

### Behavioural resistance

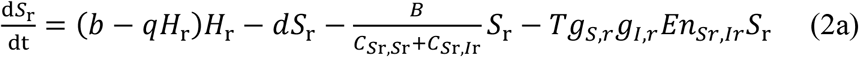

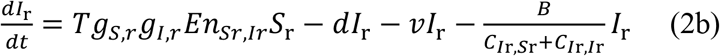

### Physiological resistance

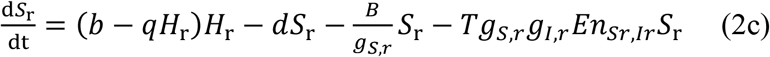

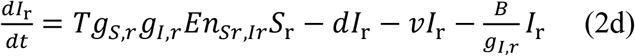

We model evolution through invasion analysis. For simplicity, we do not impose any correlation between an individual’s trait when susceptible, *g*_*S, Y*_, and its trait when infected, *g*_*I, Y*_, which may match gregariousness in the focal system [3]. The dynamics of an invading mutant follow Eq. 3:

### Behavioural resistance

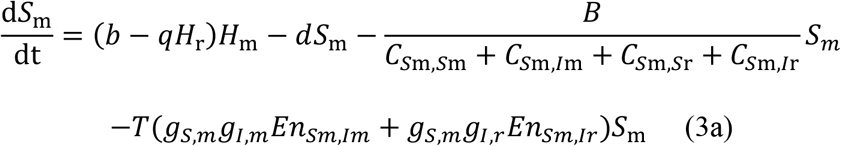

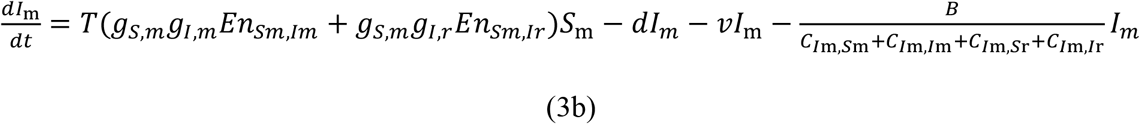

### Physiological resistance

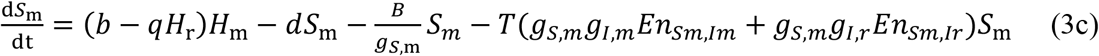

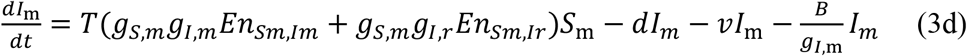

We use parameter values reasonable for the focal system. Most values were previously, empirically derived [39] but three parameter values required our judgment. First, we chose the transmissibility parameter (*T* = 0.0081) to mimic that of high predation conditions. Second, we chose encounter rate (*E* = 50) to lead to reasonable prevalence for high predation populations at intermediate resistance (*g* values around 0.5 and accounting for the fact that our model here does not include host recovery, unlike the focal system). Third, we chose *B* (0.3 for behavioural resistance and 0.006 for physiological resistance) such that hosts reaping the benefits of minimum resistance investment (*g*_*X, Y*_ = 1) experienced the same investment-dependent death rate as that of wild hosts experiencing the lowest, investment-dependent death rate in the previously published parametrization [39]. We further tested the robustness of our key results with a Latin Hypercube search of 500 parameter value sets with each parameter uniformly distributed within + 50% of the corresponding focal parameter value (except *R* which we varied uniformly from 0 to 1).

We varied the frequency of random encounters widely to capture a range of coevolutionary outcomes that may be representative for various host taxa. For clonally reproducing organisms with very limited dispersal, *R* would be approximately 0 as nearly all encounters are disproportionately with kin, even if kin have low frequency on the landscape. For other species, we can approximate qualitatively reasonable *R* values based on what would produce empirical values of mean within-group relatedness (*r* in the literature). We assume that the vast majority of an animal’s social encounters occur with members of its group. Within-group relatedness then gives the probability that another individual has the same allele from a common ancestor which we take as the portion of encounters which are with the same genotype. If a genotype has frequency *f* in the population as a whole, then *r* = *Rf*+(1-*R*) and this can be re-arranged to *R* = (*1-r*)*/*(*1-f*). When possible, we approximate between-group relatedness as *f*, the frequency of the genotype across groups; but in general, we expect *f* to be small in nature and have little impact on the calculation of *R*. For Trinidadian guppy shoals, *r* can be 0.0216 with *f* = 0.0123 giving *R* = 0.99 [27]. For various mammals, *r* can range from 0 to 0.5 [28, 29], which, assuming some small *f* such as 0.05, implies *R* from 0.52 to 1. Red-winged blackbird data can imply *R* values as low as 0.3 [(1-0.78)/(1-0.28); 26]. To capture the biological extremes, we show a spectrum of outcomes from *R* = 0 to 1.

With our empirically reasonable parameter values, we find the trait and population-level outcomes of coevolution. Numerically, we find the stable equilibrium of the resident genotype. Then we simulate a mutant at very low density and determine whether invasion fitness is positive or negative. Then we find a trait value of susceptible resistance at which evolution stops (“evolutionary singular point”). For a given rate of random encounters, *R*, we find the singular point(s) of susceptible resistance as a function of infected resistance and vice versa. An endpoint of coevolution must be an intersection of the two; further, we check the coevolutionary stability of any such intersection to determine whether populations with trait values in that neighbourhood evolve toward that intersection of trait values and stay there, making that intersection an endpoint of coevolution [55, 56]. Lastly, we calculate infection prevalence and host density for single-genotype populations to show how evolution toward a coevolutionary attractor alters population-level outcomes.

## Results

Both models agree on broad patterns of coevolution. Unsurprisingly, intersections of evolutionarily stable curves (solid blue with solid red) created coevolutionarily stable points (black dots) but if one curve was evolutionarily unstable (dotted red in Figs. 1c-e), the intersection was unstable (hence such red-blue intersections are not marked with dots in Fig. 1; see Appendix for details). There is only one coevolutionarily stable point with all random encounters (*R* = 1; Figs. 1a, b); infected hosts become selfish with coevolution favouring maximum tolerance (i.e., minimal resistance) by infected hosts (*g*_*I*, evo_ = 1) and strong resistance by susceptible hosts (low *g*_*S*, evo_; see attached code for proof that selection always favours higher *g*_*I*, evo_ for *R* = 1). Some kin association (*R* < 1; Figs. 1c-f) can lead to bistability between this selfish attractor and an altruistic attractor where infected hosts altruistically invest in resistance (low *g*_*I*, evo_) so susceptible hosts do not need to (high *g*_*S*, evo_). Perhaps surprisingly, the altruistic attractor is stable at even weak levels of kin association (e.g., *R* = 0.99, not shown), though weaker kin association does lead to smaller regions of attraction for this altruistic attractor (as seen in Fig. 1); the surprising feasibility of altruism arises due to the very costs of altruistic resistance by infected hosts. Because infected hosts pay a high cost of resistance while susceptible hosts do not, infection carries the strong, double cost of virulence *and* sharply increase resistance investment. Thus, even weak kin association creates very strong benefits of resistance investment by infected hosts (see Appendix and Fig. 1A for details) to not infect their kin and thus not shift them into expressing costly, strong resistance investment. This pattern is so strong that it can select for infected resistance to prevent spreading a mutualist; as a biologically unrealistic example to demonstrate the strength of this mechanism, we changed only the virulence of infection (*v* = −0.048) so that infected hosts live longer than susceptible hosts and kept all other aspects of the model the same. We find selection for stronger infected resistance (lower *g*_*I*, evo_) near the altruistic attractor even if resistance prevents transmission of a mutualist (we used *R* = 0.5 for both models); we found this pattern whether infected hosts reduced the spread of the mutualist behaviourally or physiologically. Our two models display these same broad patterns but their main difference is in how strongly kin association drives evolution of resistance.

**Figure 1.**
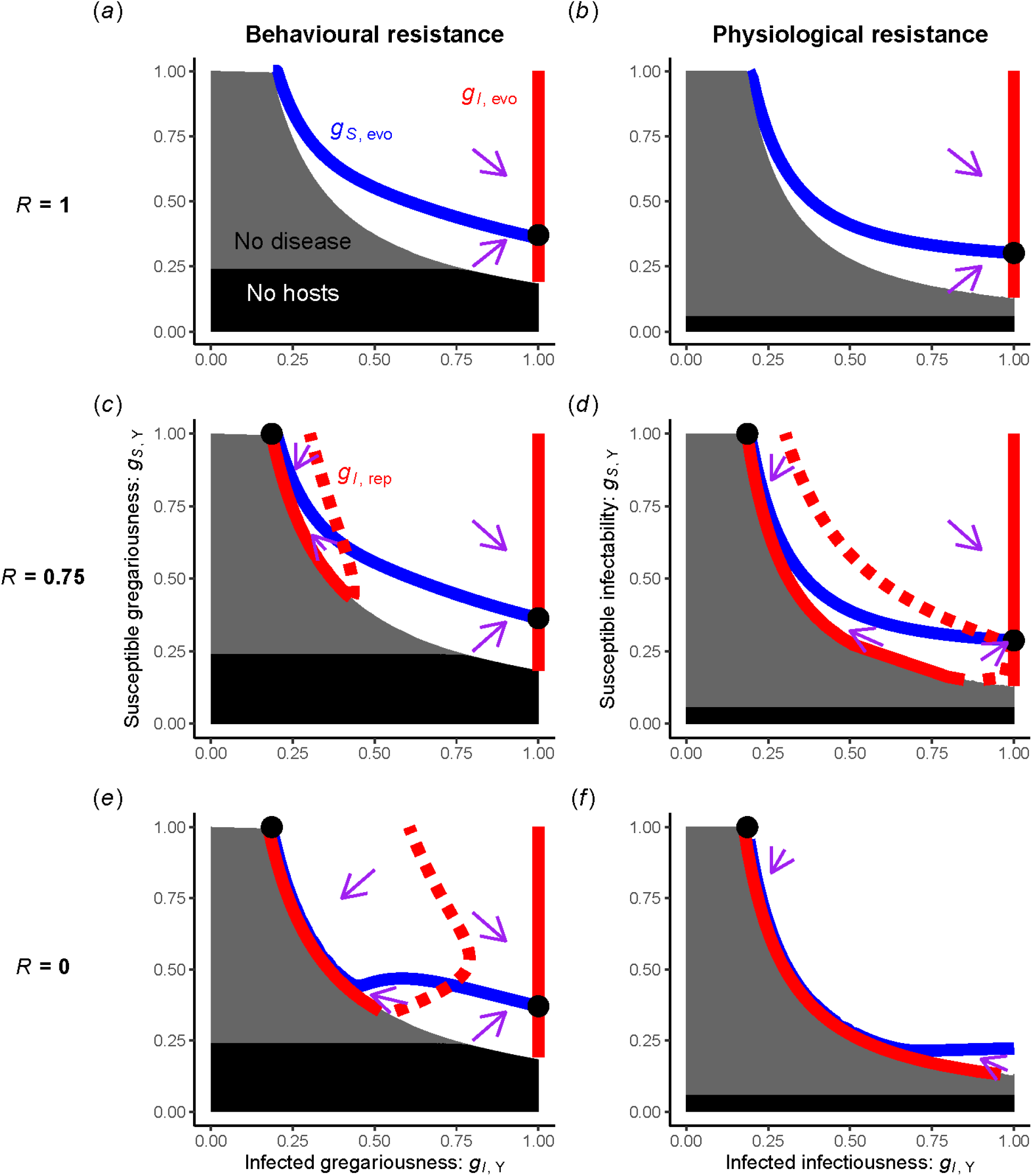
Infected hosts may selfishly spread disease through tolerance or altruistically reduce disease through resistance that protects their kin. We show the phase plane of coevolution of infected and susceptible traits. The first column shows results for the behavioural resistance model while the second shows results for the physiological resistance model. Each row shows results for a different value of the rate of random encounters, *R*. (a) If infected and/or susceptible gregariousness are too low (high resistance), the stable equilibrium may have no hosts (black rectangle) or hosts but no disease (grey rectangle). Within the parameter space with a stable, endemic equilibrium (white background), we show susceptible gregariousness that evolves at a given level of infected gregariousness (*g*_*S*, evo_, blue curve), infected gregariousness that evolves as a function of susceptible gregariousness (*g*_*I*, evo,_ red curve), the intersection of the two (black point), and the general direction of coevolution (purple arrows). (b) These results are very similar for the physiological resistance model of susceptible infectability and infected infectiousness (each the inverse of resistance). (c) If not all encounters are random (e.g., *R =* 0.75), there can be a repelling curve between two stable evolutionary outcomes of the evolution of infected gregariousness (red dotted curve, *g*_*I*, rep_), leading to different coevolutionary points. (d) The same results hold for physiological resistance except non-random encounters (lower *R*) more strongly select for resistance investment, as seen by a larger region of attraction for low infected infectiousness values. If all encounters are with kin (*R* = 0), the region of attraction becomes larger for (e) behavioural resistance or universal for (f) physiological resistance. Far left red curves shifted slightly further left in (e) and (f) for visual clarity.

The models of behavioural and physiological resistance differ in how strongly kin association drives coevolution toward the altruistic attractor. We trace this difference between the two models to terms present in the behavioural resistance model but not the physiological resistance model; these terms represent the cost of resistance investment by non-focal individuals (through fewer social contacts for the focal individual) and these terms maintain the stability of the selfish attractor even with strong kin association (see Appendix and Fig. A1). Thus, decreasing *R* leads to evolution only toward the altruistic attractor in the physiological resistance model because the costs of resistance are only borne by the individual rather than by the individual and their kin as in the behavioural resistance model.

Both models agree on the broad patterns of population-level outcomes. Host populations benefit greatly, in terms of overall density and infection prevalence, when coevolution arrives at the altruistic attractor rather than the selfish attractor. Host density is maximized when susceptible hosts have low resistance, enjoying high fitness, while infected hosts have strong resistance (Fig. 2a). This trait pairing eliminates disease (the altruistic attractor rests just on the disease-free boundary, Fig. 2b) so that all hosts enjoy the high fitness of being infection-free without strong resistance investment (because all hosts are susceptible). The selfish attractor instead leads to high prevalence of infection and low density of hosts as the host population is very infected *and* susceptible hosts are forced to maintain strong resistance investment. Results are qualitatively very similar for the physiological resistance model (Figs. 1c, d). Generally, stronger resistance investment by infected hosts improves host density; but the behavioural resistance model differs slightly by demonstrating an exception to this trend. When susceptible gregariousness is low and hosts struggle to find social contacts, higher infected gregariousness (lower resistance) can actually *increase* host density (see bottom right of Fig. 1a). This difference follows the theme of behavioural resistance’s shared cost contrasted to physiological resistance’s individual cost. Still, the two models agree strongly overall that resistance investment by infected hosts helps the host population by eliminating disease, allowing susceptible hosts to omit investment in resistance, and raising population density.

**Figure 2.**
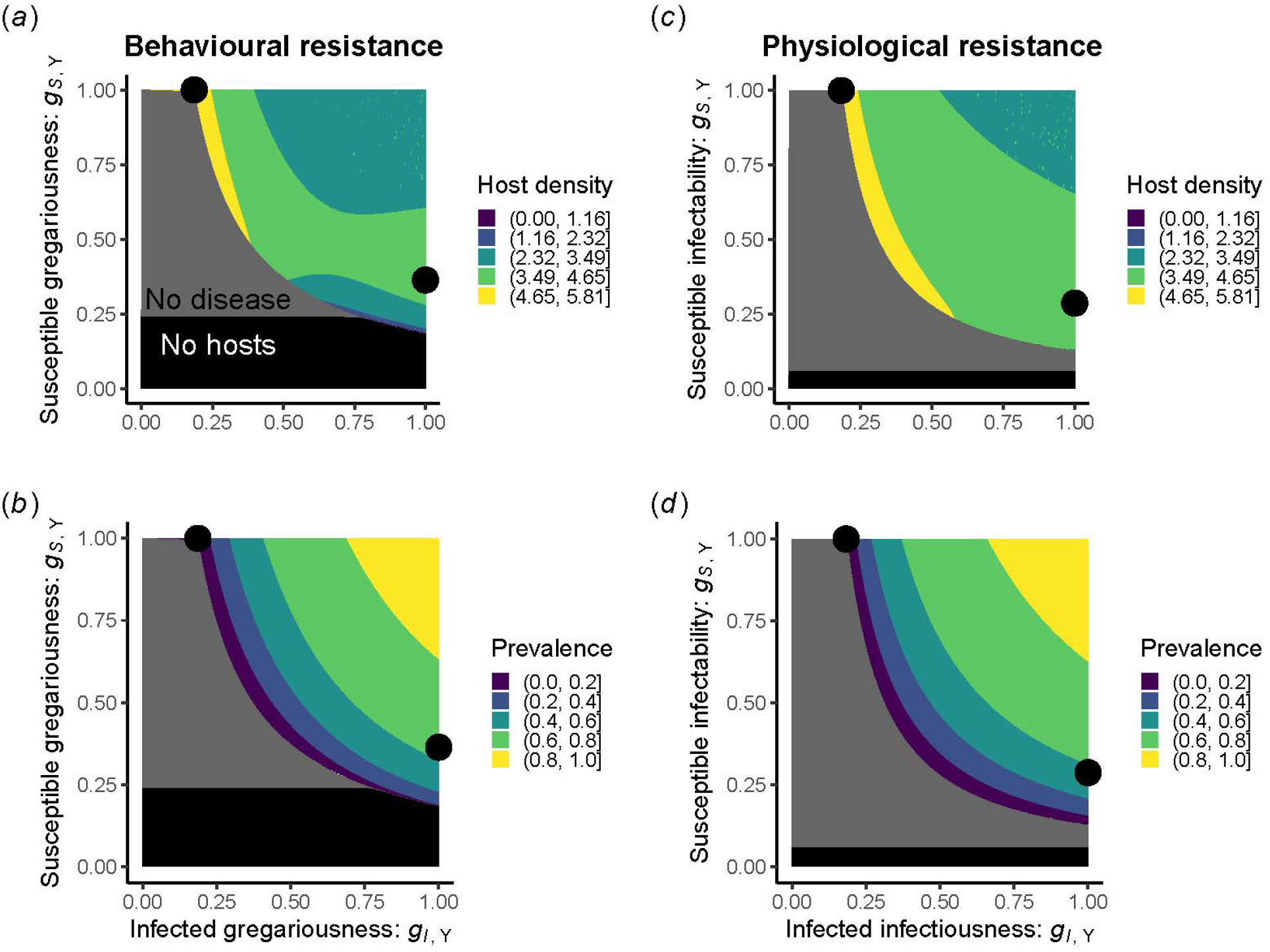
Selfish infected hosts spread disease and harm host populations while altruistic infected hosts eliminate disease and boost host populations. We show stable equilibrium host densities (top row, *H*^*^) and prevalence of infection (bottom row, *p*^*^ = *I*^*^/*H*^*^) for a population that has evolved to some pair of one infected trait value and one susceptible trait value for the behavioural resistance model (left column) and the physiological resistance model (right column). We also use black points to highlight the trait values of coevolutionary outcomes at the altruistic and selfish attractors (values shown for *R* = 0.75 but similar for other values of *R* as long as those attractors exist, see Fig. 1). The dynamics of a population with one genotype for the susceptible host trait and one genotype for the infected host trait do not depend on the value of *R* so that only the location and likelihood of coevolutionary outcomes is affected by *R*.

Our key patterns outlined here demonstrated reasonable robustness to parameter values in the parameter sets we searched (+50% for each parameter value except *R* which ranged from 0 to 1). We detected a stable altruistic attractor in 86.7% of feasible parameter sets for the behavioural resistance model and 91.5% for the physiological model. We detected a stable selfish attractor in 70.8% of feasible parameter sets for the behavioural resistance model and only 11.6% for the physiological model, reinforcing our conclusion that kin association drives selection toward the altruistic attractor more strongly in the physiological model (e.g., compare to Figs. 1e, f where only the behavioural model has a stable selfish attractor). The altruistic attractor always corresponded to higher host density and the boundary where disease goes to zero while the selfish attractor maintained disease and lower host density. Thus, our core results may have robust and broad biological relevance.

## Discussion

Considering co-evolution of infected and susceptible resistance gives a fuller picture of why selection may favour infected hosts that block transmission. Previous intuition has focused primarily on selection on either susceptible (for direct fitness) or infected resistance (for kin association), leading to a less complete picture. For example, one could consider a fixed, strong resistance investment by susceptible hosts, thus completely missing the possibility for altruism and finding that lower resistance of infected hosts increases host density (e.g., considering a horizontal slice of Figs. 1c and 2a at *g*_*S*, y_ = 0.3). Further, the costs and impacts of resistance investment by susceptible vs. infected hosts strongly influences evolution, expanding the possibility for altruism as infection shifts kin from high-fitness, low-resistance susceptible hosts to low-fitness, high-resistance infected hosts. By considering coevolution of infected and susceptible traits, our results emphasize that even weak kin association may be important in models of resistance, with large implications for prevalence and host density.

Future studies could build and improve on this work by considering how the modelling inspires new empirical work and how empirical results inspire new modelling, some of which we highlight here. Future modelling could consider whether metapopulation dynamics expand the biological relevance of the altruistic attractor. If host populations experience stochastic extinction, populations with altruistic traits are favoured for their higher host density, which is empirically associated with less extinction [57] and thus perform better in competition across a meta-population. Thus, a metapopulation model with stochastic extinction might find even more biological relevance of the altruistic attractor (note, this metapopulation reasoning does not apply to the biologically odd case of a mutualist where infected self-isolation actually reduces host density). Lastly, future models could consider various impacts of inducibility in behavioural resistance as a difference from physiological resistance. First, some hosts express their altruistic behaviour indiscriminately but some express it only for kin [29]. Second and similarly, some susceptible hosts may avoid infecteds specifically [40, 58, 59] while susceptibles may socialize with susceptibles and infecteds alike in other systems [59-62]. Outside the context of kin selection, inducibility has been found to result in some key differences between the evolution of behavioural and physiological resistance [38]; inducibility dependent on kinship and infection status could significantly change the evolution of behavioural resistance compared to physiological resistance. Thus, the intersection of resistance types and kin selection remains a fruitful area for future models.

Our modelling also suggests future empirical work. Very strong vs. weak kin association explains why infected hosts become less gregarious in social insects [18, 19] and infected bacteria kill themselves, while infected hosts become more gregarious in sticklebacks [32] or rhesus monkeys [31]. But beyond these simple cases is a broad continuum of the rate of kin association in nature. More systematic, empirical investigation in systems with moderate kin association may reveal evidence for altruistic infected resistance that blocks transmission, such as active self-isolation. For behavioural resistance, this could be found as selection for self-isolation by infected hosts that does not improve direct fitness (regardless of proximate mechanism, e.g., pathogenicity); there is some such evidence of reduced gregariousness of infected hosts outside eusocial insects [e.g., in a mouse system: 30]. However, the direct fitness effects of such reduced gregariousness should be considered as a potential confounding factor, e.g., increased activity correlated with a weaker immune response in zebra finches [22] or sick bats continued allogrooming their close kin [63]. Disentangling direct and indirect fitness effects may be worthwhile as our model motivates future investigation of self-isolation in various systems.

For physiological resistance, investment in resistance rather than tolerance could indicate altruism by infected hosts. Resistance that controls the number of parasites on an infected host reduces transmission to conspecifics [64] but may trade off with tolerance of those parasites [14-17]. Resistance may improve infection outcomes by decreasing parasite burdens or worsen infection outcomes if the sacrifice in tolerance matters more than the reduction in parasite burden. Such resistance that worsens infection outcomes may be an indication of altruism by infected hosts and could be investigated by future experiments in systems with weaker kin association than clonal bacteria [24]. All together, these modelling results emphasize that altruism by infected hosts may be more relevant than previously thought, emphasizing the importance of coevolution of infected and susceptible traits both for behavioural and physiological resistance.

