## Appendix for "Associating with kin selects against disease tolerance"

**Supplementary information for “Associating with kin selects against disease tolerance”**

**Analysis of the fitness gradient:**

We analyse how invader traits affect fitness to gain conceptual insight into the model’s results. The effect of a small change in an invader’s trait on their fitness, expressed as a partial derivative, indicates the direction of selection; because we consider mutations of small effect in our Adaptive Dynamics approach, we are interested in this quantity when the invader’s trait is similar to that of the resident (this quantity is called the fitness gradient or selection gradient). In particular, we are interested in clarifying the fitness gradient for infected host gregariousness (*g_I_*_, m_). From the fitness of an invading host (eq. A1a), we derive the expression for the fitness gradient (eq. A1b). Infected gregariousness affects fitness by affecting the prevalence of infection in the mutant genotype (∂*p*_m_/∂*g_I_*_, m_ is generally positive); increasing prevalence alters the proportion of hosts suffering parasite virulence (labelled the “Virulence” component in eq. A1b) and shifts hosts from paying the cost of susceptible immune investment to paying the cost of infected immune investment (“Infection shifts investment”). Infected gregariousness alters immune investment directly and thus affect the fitness of susceptible hosts (“Investment affects *S*”) and infected hosts (“Investment affects *I*”). Similar components can be derived for the physiological immunity model (eqs. A1c, A1d) except that changing immune investment of infected hosts does not affect the investment cost-associated part of the fitness of susceptible hosts (i.e., there is no “Investment affects *S*” component in eq. A1d).

*Behavioural immunity:*

$\frac{1}{H_{m}}\frac{dH_{m}}{\mathrm{dt}}=b-qH_{r}-d-\underset{Immune investment}{\underbrace{\left( \frac{B}{C_{Sm,HT}}\left( 1-p_{m} \right)+\frac{B}{C_{Im,HT}}p_{m} \right)}}-\underset{\mathrm{Virulence}}{\underbrace{vp_{m}}}$	(A1a)

$\frac{\partial}{\partial g_{I, m}}\frac{1}{H_{m}}\frac{dH_{m}}{\mathrm{dt}}=-\underset{\mathrm{Virulence}}{\underbrace{v\frac{\partial p_{m}}{\partial g_{I, m}}}}+\underset{Infection shifts investment}{\underbrace{\left( \frac{B}{C_{Sm,HT}}-\frac{B}{C_{Im,HT}} \right)\frac{\partial p_{m}}{\partial g_{I, m}}}}+\underset{Investment affects S}{\underbrace{\frac{B(1-p_{m})}{(C_{Sm,HT})^{2}}\frac{\partial C_{Sm,HT}}{\partial g_{I, m}}}}+\underset{Investment affects I}{\underbrace{\frac{Bp_{m}}{(C_{Im,HT})^{2}}\frac{\partial C_{Im,HT}}{\partial g_{I, m}}}}$	(A1b)

*Physiological immunity:*

$\frac{1}{H_{m}}\frac{dH_{m}}{\mathrm{dt}}=b-qH_{r}-d-\underset{Immune investment}{\underbrace{\left( \frac{B}{g_{S,m}}\left( 1-p_{m} \right)+\frac{B}{gI,m}p_{m} \right)}}-\underset{\mathrm{Virulence}}{\underbrace{vp_{m}}}$	(A1c)

$\frac{\partial}{\partial g_{I, m}}\frac{1}{H_{m}}\frac{dH_{m}}{\mathrm{dt}}=-\underset{\mathrm{Virulence}}{\underbrace{v\frac{\partial p_{m}}{\partial g_{I, m}}}}+\underset{Infection shifts investment}{\underbrace{\left( \frac{B}{gS,m}-\frac{B}{gI,m} \right)\frac{\partial p_{m}}{\partial g_{I, m}}}}+\underset{Investment affects I}{\underbrace{\frac{Bp_{m}}{{(g}_{I,m})^{2}}}}$	(A1d)

By examining how immune investment affects host fitness, we can see how only the behavioural model captures the costs of immune investment by one individual for other individuals. In the physiological immunity model, immune investment by infected hosts only affects the investment cost-associated part of fitness for a given focal host (“Investment affects *I*” in eq. A1d). In the behavioural immunity model, infected investment alters the investment-cost associated part of the fitness of susceptible hosts by altering the total contact rate enjoyed by susceptible hosts of the mutant genotype (*C_S_*_m,_ *_HT_* in “Investment affects *S*” in eq. A1b). This total contact rate depends on the gregariousness of focal individuals and non-focal individuals (eq. A2a). Infected gregariousness alters the contact rate for susceptible hosts directly when an infected host serves as the non-focal individual (“N: *I* as non-focal” in eq. A2b with the N label denoting that this arises from the investment of non-focal individuals). Infected gregariousness also shifts how many of a focal, susceptible host’s non-random encounters are with susceptible hosts expressing susceptible gregariousness (*g_S,_* _m_) vs. infected hosts expressing infected gregariousness (*g_I_*_, m_; feeding into the “N: Composition of encounters change” part of eq. A2b). Because both of these factors quantify the cost of immune investment in terms of the traits of non-focal individuals, we consider both to be part of the “cost of non-focal immune investment” that arises here and drives the ”Investment affects S” term in eq. A1b. We use a similar derivation for the contact rate enjoyed by infected hosts (eq. A2c) and find similar terms (driving part of “Investment affects *I*” in eq. A1b) plus one more; there is a term capturing the effect of infected gregariousness on contact rate acting through the trait of the focal individual (“F: *I* as focal” in eq. A2d with F for focal, which also drives “Investment affects *I*”; see eq. A2e for how these terms factor into ). Thus, both models have a component that is the “cost of focal immune investment” while only the behavioural model has a term that is the “cost of non-focal immune investment”. The behaviour of the shared fitness terms between the two models helps us understand similarities in model outcomes while the behaviour of this extra, non-focal term helps us understand differences in model outcomes (Fig. A1).

$C_{Sm,HT}=ERg_{S, m}g_{S,r}(1-p_{r})+ERg_{S, m}g_{I,r}p_{r}+E(1-R)g_{S, m}g_{S,m}(1-p_{m})+E(1-R)g_{S, m}g_{I,m}p_{m}$	(A2a)

$\frac{\partial C_{Sm,HT}}{\partial g_{I, m}}=\underset{N: I as non-focal}{E\underbrace{\left( 1-R \right)g_{S, m}p_{m}}}+\underset{N: Composition of encounters change}{E\underbrace{\left( 1-R \right)g_{S, m}\frac{\partial p_{m}}{\partial g_{I,m}}\left( g_{I,m}-g_{S,m} \right)}}$	(A2b)

$C_{Im,HT}=ERg_{I, m}g_{S,r}(1-p_{r})+ERg_{I, m}g_{I,r}p_{r}+E(1-R)g_{I, m}g_{S,m}(1-p_{m})+E(1-R)g_{I, m}g_{I,m}p_{m}$	(A2c)

$\frac{\partial C_{Im,HT}}{\partial g_{I, m}}=\underset{F: I as focal}{\underbrace{ERg_{S,r}\left( 1-p_{r} \right)+ERg_{I,r}p_{r}+E\left( 1-R \right)g_{S,m}\left( 1-p_{m} \right)+E\left( 1-R \right)g_{I, m}p_{m}}}+\underset{N: I as non-focal}{\underbrace{E\left( 1-R \right)g_{I, m}p_{m}}}+\underset{N: Composition of encounters change}{E\underbrace{\left( 1-R \right)g_{I, m}\frac{\partial p_{m}}{\partial g_{I,m}}\left( g_{I,m}-g_{S,m} \right)}}$	(A2d)

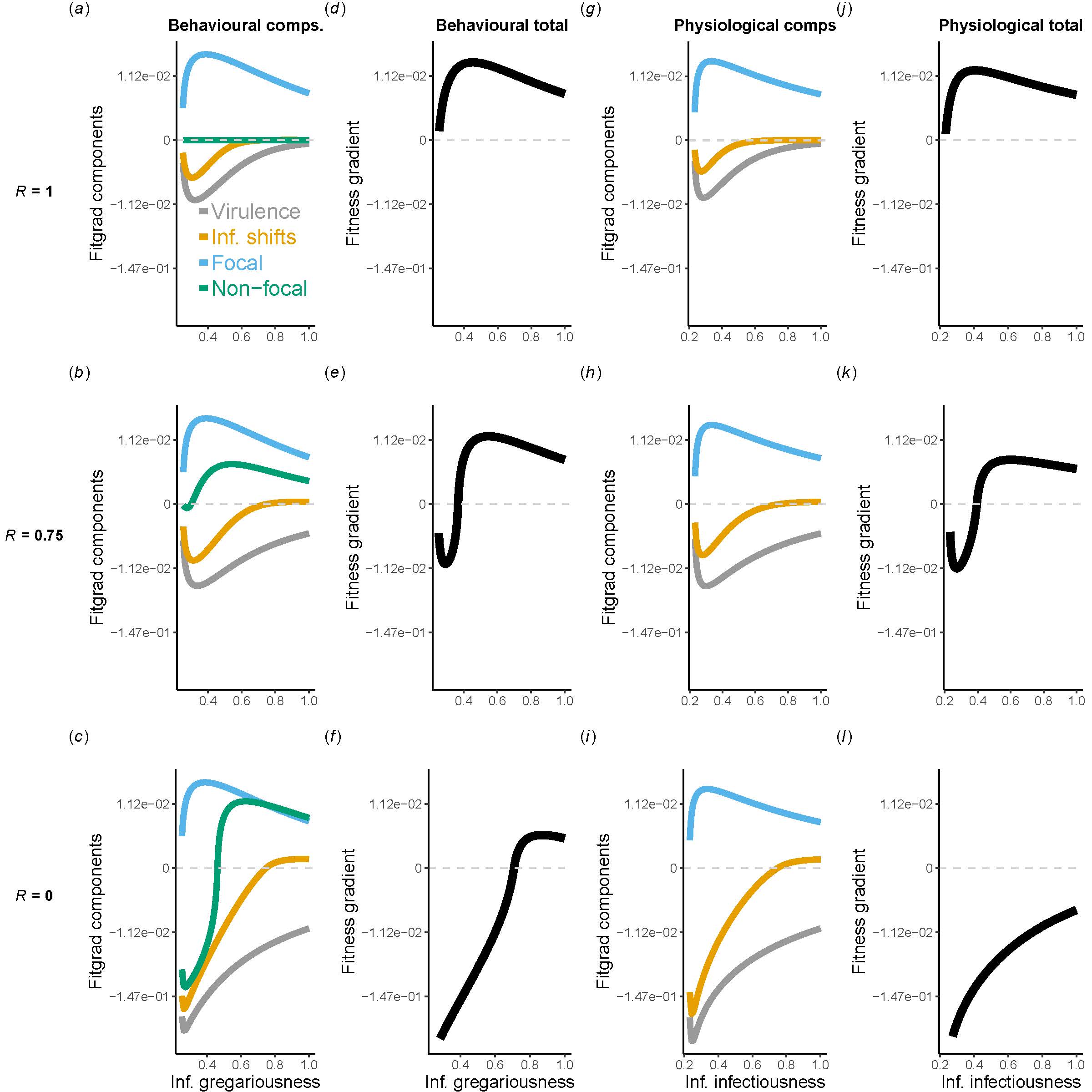

**Figure A1. Components of the fitness gradient help understand coevolutionary results.** We plot components of the fitness gradient and the total fitness gradient across levels of *R* (ranging from no kin selection, *R* = 1, to very strong, *R* = 0), for both models, across infected investment, and for a fixed, low level of susceptible investment (*g_S, Y_* = 0.747). The components of the fitness gradient are virulence (grey), infection shifting investment by shifting host classes (gold), the cost of focal (blue) immune investment, and the fitness cost of non-focal immune investment (green). Positive values indicate selection for higher gregariousness for the behavioural model or higher infectiousness for the physiological model (both correspond to more transmission and lower immune investment). Note from tick labels that the y-axis is scaled non-linearly to emphasize values near zero as these small magnitude values can critically determine the sign of the fitness gradient.

Examining the components of the fitness gradient clarifies why both models show stability of the “altruistic” attractor given some kin selection and why the “selfish” attractor maintains stability better in the behavioural immunity model. With no kin selection (*R* = 1, Fig. A1a), virulence and infection shifting investment are relatively small factors; this is because kin-kin contact is negligible for rare mutants so higher mutant infected gregariousness (*g_I,_* _m_) only weakly increases prevalence by increasing how long infected hosts survive, not by increasing transmission. So the benefits of decreasing focal immune investment are larger. Stronger kin selection (moving to *R* = 0.75 in Fig. A1b and *R* = 0 in Fig. A1c) has little effect on this focal immune investment term but strengthens all other terms including the cost of non-focal immune investment as kin-kin contacts become non-negligible. Totalling the components of the fitness gradient, the fitness gradient is always positive, selecting for higher infected gregariousness, with no kin selection (Fig. A1d; see attached code for proof). With stronger and stronger kin selection, there is a larger and larger region of low gregariousness values that lead to selection for lower gregariousness (negative fitness gradient) and a smaller region of high gregariousness values that lead to selection for higher gregariousness (positive fitness gradient; Figs. A1e, A1f). Results are similar for the physiological immunity model (Figs. A1g-l) except the non-focal immune investment is absent.

In both models, virulence and infection shifting immune investment maintain the stability of the “altruistic” attractor where infected hosts invest strongly in immunity to eliminate disease (low *g_I_*_, evo_) while susceptible hosts do not need to invest in immunity (*g_S_*_, evo_ = 1; see Fig. 1). When mutant infected hosts transmit disease more (*g_I_*_, m_), the mutant population suffers more infection (especially for *R* < 1 where mutants disproportionately encounter kin). Thus, more mutant hosts bear the cost of infection (virulence term selects for lower *g_I_*_, m_). Less obviously, infection shifts immune investment in an unfavourable manner near the “altruistic” attractor. In the neighbourhood of this attractor, infected hosts pay a high fitness cost of immune investment (low *g_I,_* _m_) while susceptible hosts pay a minimal fitness cost (*g_S_*_, m_ = 1), making the infection shifts investment a strong factor selecting for lower *g_I_*_, evo_ to avoid shifting hosts into the extra-unfavourable infected class. This logic applies in both models (Fig. A1a).

But uniquely to the behavioural model, the cost of non-focal immune investment maintains stability of the “selfish” attractor. Near the “altruistic” attractor, it can be seen that the cost of non-focal immune investment must select for lower infected gregariousness (*p*_m_ ≈ 0 and *g_S,_* _m_ *> g_I,_* _m_ so “*I* as non-focal” ≈ 0 and “Composition of encounters change” < 0). Near the “selfish” attractor, prevalence is high and *g_I,_* _m_ *> g_S,_* _m_ so the cost of non-focal immune investment is a positive term, selecting for higher infected gregariousness (also see Fig. A1a-c). The fitness gradient terms for the two models are quite similar except for this cost of non-focal immune investment term in the behavioural immunity model, which maintains stability of the “selfish” attractor (compare Fig. A1c to Fig. A1i to see how green curve contributes to positive total at high infected gregariousness in Fig. A1f while total remains negative in Fig. A1l).

**Analysis of coevolutionary stability**

Because we assume no correlation between the trait of infected hosts (*g_I,_* _r_ for some resident r) and the trait of susceptible hosts (*g_S_*_, r_ for some resident), the conditions for coevolutionary convergence stability are quite simple. Followomg Leimar 2009, the mutational matrix ***K*** can be any diagonal matrix with positive diagonal elements. Coevolutionary convergence stability also depends on the Jacobian of the selection gradient, ***J***, in eq. A3:

$\boldsymbol{J}\mathbf{=}\left[ \begin{matrix} \frac{d}{dg_{S, r}}\left( \frac{\partial}{\partial g_{S, m}}\frac{1}{H_{m}}\frac{dH_{m}}{\mathrm{dt}} \right)|_{g_{S, m}=g_{S, r}} & \frac{d}{dg_{I, r}}\left( \frac{\partial}{\partial g_{S, m}}\frac{1}{H_{m}}\frac{dH_{m}}{\mathrm{dt}} \right)|_{g_{S, m}=g_{S, r}} \\ \frac{d}{dg_{S, r}}\left( \frac{\partial}{\partial g_{I, m}}\frac{1}{H_{m}}\frac{dH_{m}}{\mathrm{dt}} \right)|_{g_{I, m}=g_{I, r}} & \frac{d}{dg_{I, r}}\left( \frac{\partial}{\partial g_{I, m}}\frac{1}{H_{m}}\frac{dH_{m}}{\mathrm{dt}} \right)|_{g_{I, m}=g_{I, r}} \end{matrix} \right]$(A3)

An intersection of evolutionary singular points for *g_S_*_, evo_ and *g_I,_* _evo_ will be convergence stable if and only if det(***J***) > 0 (determines sign of determinant of ***AJ***) and ***A*_11_*J*_11_+*A*_22_*J*_22_** < 0 (trace of ***AJ***). We find that the intersections of a convergence stable *g_S_*_, evo_ curve with a convergence stable *g_I_*_, evo_ curve satisfy these conditions (see attached code) while the intersections of a convergence stable *gS,* evo curve with a convergence unstable *gI, evo* curve did not satisfy the determinant condition. We also confirmed the evolutionary stability (i.e., uninvasibility) of our convergence stable intersections. Thus, these intersections represent endpoints of coevolution toward which similar populations will evolve and at which the population will remain.
